# Meiotic sister chromatid exchanges are rare in *C. elegans*

**DOI:** 10.1101/2020.07.22.216614

**Authors:** David E. Almanzar, Spencer G. Gordon, Ofer Rog

## Abstract

Sexual reproduction shuffles the parental genomes to generate new genetic combinations. To achieve that, the genome is subjected to numerous double-strand breaks, the repair of which involves two crucial decisions: repair pathway and repair template. Use of crossover pathways with the homologous chromosome as template exchanges genetic information and directs chromosome segregation. Crossover repair, however, can compromise the integrity of the repair template and is therefore tightly regulated. The extent to which crossover pathways are used during sister-directed repair is unclear, because the identical sister chromatids are difficult to distinguish. Nonetheless, indirect assays have led to the suggestion that inter-sister crossovers, or sister chromatid exchanges (SCEs), are quite common. Here we devised a technique to directly score physiological SCEs in the *C. elegans* germline using selective sister chromatid labeling with the thymidine analog 5-ethynyl-2’-deoxyuridine (EdU). Surprisingly, we find SCEs to be rare in meiosis, accounting for <2% of repair events. SCEs remain rare even when the homologous chromosome is unavailable, indicating that almost all sister-directed repair is channeled into noncrossover pathways. We identify two mechanisms that limit SCEs. First, sister-directed repair intermediates are efficiently inhibited by the RecQ helicase BLM^HIM-6^. Second, the Synaptonemal Complex–a conserved interface that promotes crossover repair– localizes between the homologous chromosomes and not the sister chromatids. Our data suggest that in *C. elegans* crossover pathways are only used to generate the single necessary link between the homologous chromosomes. Almost all other breaks, regardless of which repair template they use, are repaired by noncrossover pathways.

## Results and Discussion

### Selective labeling of a single sister chromatid using EdU

During meiosis, chromosomes receive multiple double-strand breaks (DSBs), but undergo only a very limited number of inter-homolog exchanges, or crossovers [1–4]. These repair events manifest as chiasmata, the obligate physical links that enable correct segregation of the homologous chromosomes (homologs). The remaining breaks are repaired by alternative methods, including repair using the homolog without exchange of flanking sequences (noncrossover), and repair using the sister chromatid as a template, forming both crossovers and noncrossovers. While meiotic repair is biased toward the homolog [5], the prevalence and regulation of sister-directed repair is poorly understood, largely due to challenges in distinguishing the identical sisters and consequently, the resulting repair products. Inter-sister crossover products, or sister chromatid exchanges (SCEs), unlike inter-sister noncrossover products, can be directly identified using nucleotide analogs [6–9]. However, these assays have not been extensively applied to modern model systems where meiotic recombination is well characterized and SCE etiology can be ascertained. To address this, we took advantage of the cytological and genetic amenability of *C. elegans*, and developed a way to selectively label individual sister chromatids. This allowed us to quantify SCEs and to study SCE regulation.

Inspired by the classical experiments of Meselson and Stahl [10], we devised a pulse/chase method that results in only one sister containing the thymidine analog 5-ethynyl-2’-deoxyuridine (EdU). The distal end of the *C. elegans* gonad is populated by mitotically dividing germline nuclei that eventually enter into meiosis and progress through meiotic prophase as they travel through the gonad (Fig. 1A). We exposed worms to a short pulse of EdU via soaking, allowing it to be incorporated into newly replicated DNA for one mitotic S-phase (Fig. 1B and S1; [2,11,12]). Chase times were then optimized such that labeled nuclei would enter meiosis, undergo premeiotic S-phase without EdU, and complete meiotic prophase, resulting in one of the strands of one of the two sisters containing EdU. (An additional mitotic cycle without EdU prior to meiotic entry would result in a combination of one or no EdU-containing sisters in each chromosome. See Methods for more details.) The azide group on EdU can be labeled with an organic dye using ‘click’ chemistry [13], which is easily compatible with other labeling modalities and does not require DNA denaturation, unlike antibody labeling of 5-bromo-2’-deoxyuridine (BrdU; [14]).

**Figure 1:**
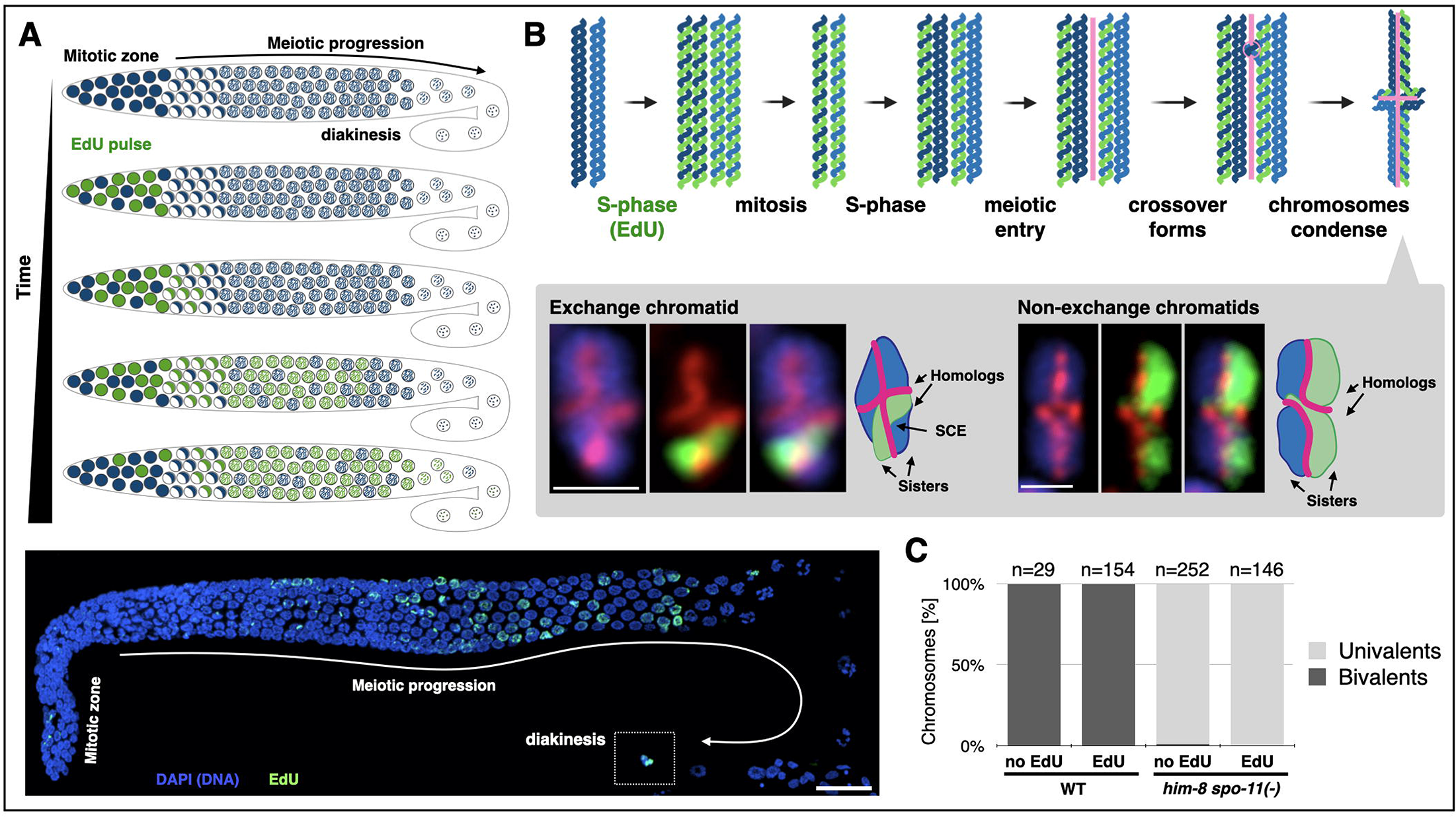
A new method to visualize meiotic SCEs using selective labeling with EdU. A. Overview of pulse/chase experiment. Nuclei undergoing DNA replication in the mitotic zone of the gonad incorporate EdU (green). After removal of EdU, the labeled nuclei continue dividing asynchronously until meiotic entry (crescent-shaped nuclei). The labeled nuclei progress through meiosis as they move through the gonad, and are remodeled in diakinesis into condensed structures that allow visualization of individualized sisters. Bottom, an image of a wildtype *C. elegans* gonad labeled with EdU. A labeled nucleus in diakinesis is marked with a dashed box. Note the asynchronous distribution of the labeled nuclei. Blue, DNA; Green, EdU. B. Schematic illustrating single sister EdU labeling. Chromosomes undergo DNA replication in the presence of EdU (green), resulting in one EdU-containing strand in each sister. Following pre-meiotic DNA replication without EdU, labeling is reduced to one strand of one sister per homolog pair. In meiotic prophase, the meiotic chromosome axis (pink) appears at the interface between the homologs. When visualized as a condensed bivalent in diakinesis, sister chromatids are clearly resolvable, with the axis now delineating the interface between the sisters. Inset, a confocal image of bivalents harboring or lacking an exchange (left and right, respectively) with interpretative diagrams to the right. The nucleus harboring the exchange bivalent underwent an extra round of DNA replication without EdU before entering meiosis, resulting in labeling of only one of the four sisters. Blue, DNA (DAPI); red, axis (anti-HTP-3 antibodies); green, EdU. Scale bars = 1μm. C. EdU incorporation does not cause measurable DNA damage or affect bivalent formation. Analysis of bivalent formation in wildtype and *him-8 spo-11(−)* worms, which lack meiotic DSBs. (*him-8* is present in this background due to genetic linkage with *spo-11*, but this does not impact the result.) DSB formation in *him-8 spo-11(−)* animals would result in chiasmata and bivalent formation; no bivalents were observed among the EdU labeled chromosomes. The presence of univalents in wildtype animals would have indicated failure to form chiasmata; no such incidence were observed in the presence or absence of EdU. N values indicate the number of chromosomes counted per genotype. See Fig. S2 for further analysis.

At the end of meiotic prophase (diakinesis) chromosomes condense and sister chromatids individualize. The paired homologs are linked together by chiasmata, forming a cruciform-like bivalent, with each sister on either side of the meiotic chromosome axis (stained here with antibodies against the axis protein HTP-3 [15]; Fig. 1B, inset). By combining EdU labeling with DAPI staining of bulk chromatin we directly visualized single sister chromatids. SCEs manifest at this stage as an EdU signal traversing the axis, analogous to the so-called “harlequin chromosomes” ([6,7]; Fig. 1B, inset).

We wanted to address the potential impact of our experimental approach on genome integrity and DNA repair. In worms lacking endogenous meiotic DSBs even a single break is sufficient for chiasma formation [2,3], providing us with a sensitive readout for DNA damage. In animals lacking SPO-11, the enzyme that catalyzes meiotic DSBs [16], the homologs do not form a chiasma, and therefore condense as two unassociated oval univalents, with the two sisters on either side of the axis. EdU incorporation did not significantly induce chiasmata in this background (Fig. 1C and Fig. S2), indicating that EdU substitution does not induce DNA breaks in meiotic prophase. Wildtype animals labeled with EdU form chiasmata as normal (Fig. 1C and Fig. S2), indicating no significant perturbation of the crossover repair pathway. In these animals we also did not observe any chromosome fragmentation (n=183 EdU labeled chromosomes) suggesting DNA repair was functional. We conclude that our EdU labeling approach can reliably capture physiological sister-directed repair, overcoming challenges reported for BrdU labeling [17,18].

### SCEs are a rare outcome of meiotic DNA repair

When we applied our single-sister labeling approach to wildtype animals, we observed only 4% of chromatids harboring exchanges, indicating a low level of SCEs (2/49 chromosomes; Fig. 2A; all SCE counts are summarized in Table S1; statistically significant pairwise comparisons are noted). This result might stem from a very strong bias for homolog-directed repair, which in *C. elegans* can only occur between already paired homologs. To investigate whether SCEs are elevated when only the sister chromatid is available as a template for repair, we used *him-8* animals where the X chromosome fails to pair with its homolog and engage in homolog-directed repair, but programmed DSBs are formed and are successfully repaired [19]. Even in this condition we observed very few exchange chromatids (1/24 univalents; Fig. 2A-B), indicating that competition with the homolog is not the reason for the paucity of SCEs. Alternative pathways that do not use a repair template, such as canonical nonhomologous end-joining (c-NHEJ), single-strand annealing (SSA) or theta-mediated end-joining (TMEJ), are thought to be rarely used during *C. elegans* meiosis [20–23]. Nonetheless, we directly examined the role of c-NHEJ by eliminating DNA ligase IV (*lig-4*; [24]). While breaks form and are efficiently repaired in *lig-4; him-8* worms, we observed very few SCEs on both univalents and bivalents (0/28 and 1/55, respectively; Fig. 2A), indicating that the low rate of SCEs in *him-8* animals does not stem from breaks being channeled to c-NHEJ. Taken together, these data indicate that SCEs are a rare outcome of meiotic DSB repair, whether the homolog is available, and that sister-directed repair during meiosis almost always results in noncrossovers.

**Figure 2:**
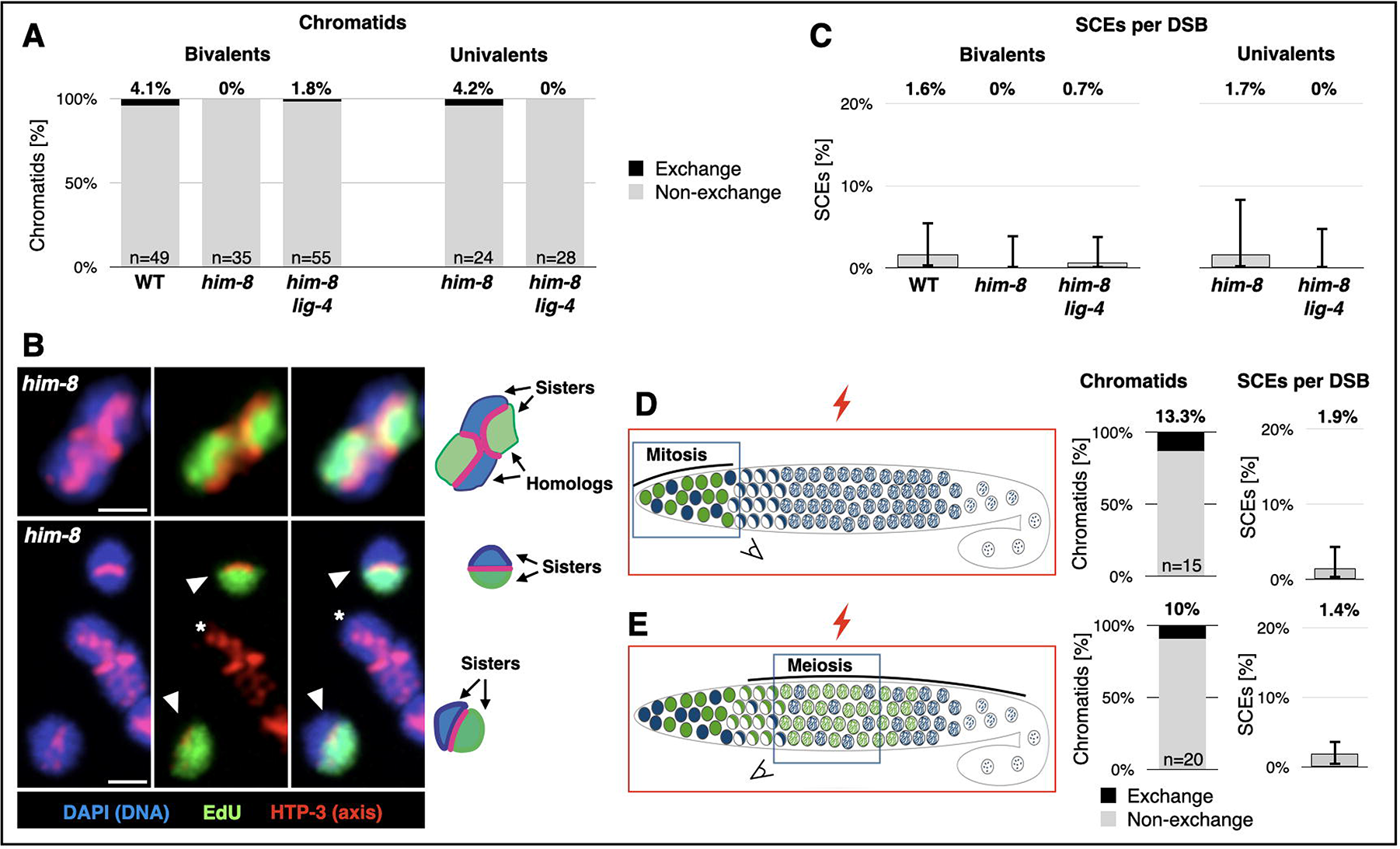
SCEs are rare outcome of meiotic DSB repair. A. Exchange chromatids are rare, regardless of the availability of the homolog. Exchange and non-exchange chromatids on bivalent and univalent chromosomes in wildtype worms, in worms deficient for X chromosome pairing (*him-8*) and in worms defective for both X chromosome pairing and non-homologous end-joining (*him-8; lig-4*). N values indicate the number of chromatids counted for each genotype. Results were not significant for all pairwise comparisons with wildtype chromosomes (p>.05, Pearson’s chi-squared test). B. Representative cytology of bivalents and univalents from *him-8* animals. Top, a single sister of each homolog pair labeled with EdU, with no exchanges in either of the long arms. Bottom, two unpaired univalent X chromosomes (arrowheads) in *him-8* animals, each with a single sister chromatid labeled with EdU, and no exchanges. An autosome bivalent (asterisk) is not labeled due to the different replication time of the X chromosome and the autosomes [64]. Red, axis (anti-HTP-3 antibodies); green, EdU, Blue, DNA (DAPI). Interpretive diagrams are shown on the right. Scale bars = 1μm. C. SCEs are a rare outcome of meiotic DSB repair. Rates of SCE formation per DSB calculated using an estimate of 5 DSBs per homolog pair (2.5 per chromosome). Numbers represent the average rate a DSB resulted in an SCE. Error bars represent 95% exact binomial confidence intervals. D. Direct quantification reveals that SCEs are a rare outcome of exogenous DSBs. Left, cartoon of irradiation experiment. *him-8 spo-11(−)* worms that are deficient for endogenous DSBs are labeled with EdU and exposed to 5000 rads X-ray irradiation (an average of 7 DSBs per chromosome). While the whole worms were subject to irradiation (red box), the nuclei in the mitotic zone (D) and meiotic prophase (E) reached diakinesis at time of dissection (indicated by blue squares). Middle, observed exchange chromatids. N values indicate the number of chromatids counted for each genotype. Right, rate of SCEs as a fraction of total DSB repair events. Error bars represent 95% exact binomial confidence intervals.

The prevalence of exchange chromatids counted above (Fig. 2A) overestimates the rate of SCE formation, since each chromosome is subjected to multiple DSBs. However, the number of meiotic DSBs in *C. elegans* remains controversial, with results from different assays ranging from 2 to more than 8 per homolog pair [1,2,25,26]. Therefore, in the experiments above we could only estimate the rate at which a DSB repair event results in an SCE, revealing rates of 1-2% SCEs per DSB in the various conditions we tested (Fig. 2C; calculated using an estimate of 5 DSBs per homolog pair, or 2.5 DSBs per chromosome; 95% exact binomial confidence intervals are shown). In the case of *him-8* and *lig-4; him-8* animals this is likely a conservative estimate, since these mutants seem to incur additional breaks compared with wildtype animals [27,28]. To allow for direct quantification of the rate of SCE formation we induced a known number of DSBs in mitotic or meiotic nuclei using X-ray irradiation in *him-8 spo-11(−)* animals that lack endogenous breaks (Fig. 2D-E). We irradiated the entire gonad at different times following EdU labeling, and identified the cell cycle stage at the time of DSB induction using chromosome morphology: chromosomes subjected to DSBs in meiosis subsequently formed bivalents connected by chiasmata, whereas chromosomes subjected to DSBs in the mitotic zone condensed as univalents [16]. Upon induction of an average of 7 DSBs/chromosome (see Methods for details), mitotic and meiotic nuclei harbored 13.3% and 10% exchange chromatids, respectively. This rate translates to 1.9% and 1.4% of DSBs repaired as SCEs (Fig. 2D-E). The similar low rate of meiotic SCEs induced by irradiation and the estimated rate for endogenous DSBs suggests that neither the nature of X-ray induced breaks nor their genomic location play a significant role in determining repair outcomes. Furthermore, our analysis of mitotic chromosomes and of unpaired homologs in meiosis suggests that the low rate of SCEs is not a result of interaction or competition with the homolog.

Investigations of inter-sister repair have typically measured imperfect repair products between repeated elements, where the homolog is not available and only sister-directed repair can occur (e.g., [29]). These studies, like our data, indicate a disfavoring of crossover *versus* noncrossover outcomes during meiotic inter-sister repair. This conclusion receives further substantiation in the accompanying work in *C. elegans* by Toraason, Libuda and coworkers.

Our labeling approach also enabled us to measure the frequency of SCEs in unperturbed meiosis, revealing that 4% of chromatids harbor an exchange, and 1.4-1.7% of DSBs are repaired as SCEs. Basal SCE rates of 10-20% were reported on both the Drosophila X chromosome and on chromosome III in budding yeast [30–33]. However, these measurements were made in cells with altered karyotype, relying on topological entrapments caused by SCEs in circular (or “ring”) chromosomes. In yeast, repair intermediates could be identified using physical assays that rely on obtaining synchronous cultures where specific loci – “hotspots” – undergo meiotic DSBs at very high rates. Quantification of inter-homolog and inter-sister intermediates unique to crossover pathways, such as double Holliday Junctions, suggested that inter-sister events comprise ~20% of all joint molecules in budding yeast, and as much as 75% in fission yeast [5,34–37]. These rates translate to 8-25% of all sister-directed events, and 2.5-9% of all DSBs, resulting in SCEs (low and high estimates are calculated based on [35]). The higher rates compared to our analysis of SCEs in *C. elegans* could be explained by evolutionary divergence of repair pathways and their regulation, or as a consequence of the very “hot” hotspots that were analyzed, which do not represent the rest of the yeast genome (incidentally, *C. elegans* lacks strong meiotic hotpots [38,39]). Alternatively, the resolution of inter-sister repair intermediates may be heavily biased toward a noncrossover fate, so that SCEs in yeast meiosis are quite rare, as we find in worms.

### The RecQ helicase BLM^HIM-6^ antagonizes meiotic SCEs

The scarcity of meiotic SCEs in worms could be explained by a complete lack of sister engagement by the crossover repair machinery, or by efficient destabilization of sister-directed crossover intermediates. RecQ helicases play multiple roles during meiosis and mitosis, including disassembling of repair intermediates ([40–43]; reviewed in [44]), compelling us to assess the prevalence of meiotic SCEs in worms lacking the RecQ helicase BLM^HIM-6^ (designated as *him-6* worms). While eliminating BLM^HIM-6^ does not affect homolog pairing or the formation of inter-homolog crossover precursors [40], meiotic SCEs were greatly elevated in *him-6* animals (26/60 chromosomes; Fig. 3A-B). The majority of SCEs likely result from meiotic events, since SCE levels were significantly lower in *him-6 spo-11* worms that lack meiotic DSBs (6/49 chromosomes; Fig. 3B-C). These residual SCEs likely reflect problems during DNA replication (Fig. 3D), as was reported for RecQ deficiencies in worms and other systems ([45], reviewed in [44]). This analysis suggests that sister chromatids are engaged in crossover repair, but these repair intermediates are efficiently destabilized by BLM^HIM-6^, explaining why so few SCEs form in meiosis. Since the rate of SCE formation is still limited in *him-6* animals (<17% of DSBs repaired as SCEs, calculated as in Fig. 2C), additional mechanisms must also act to prevent meiotic SCEs. Application of our assay to genetic perturbations that implicated other molecules in the regulation of sister-directed repair is likely to shed light on these mechanisms ([46–49] and accompanying manuscript by Thoraason, Libuda and coworkers).

**Figure 3:**
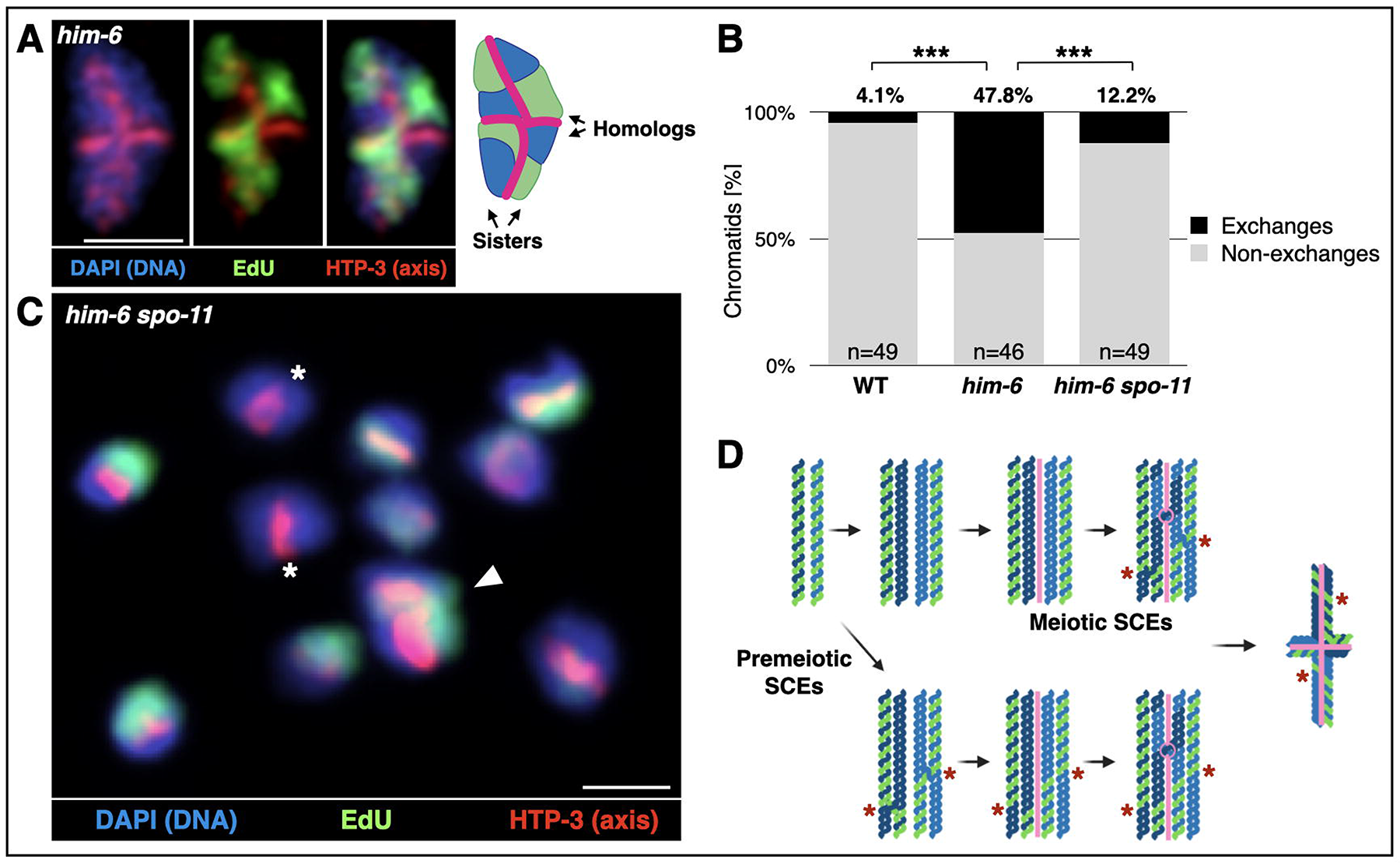
SCEs are elevated in worms lacking the RecQ helicase BLM^HIM-6^. A. Multiple SCEs are visible in an immunofluorescence image of a bivalent from a *him-6* worm. Note that each sister pair underwent an SCE. Red, axis (anti-HTP-3 antibodies); green, EdU, Blue, DNA (DAPI). Interpretive diagram shown to the right. Scale bar = 1μm. B. Elevated levels of meiotic SCEs in *him-6* worms. Frequency of observed exchange chromatids in wildtype, *him-6* and *him-6 spo-11* worms. Note the much lower level of exchanges in *him-6 spo-11 versus him-6* worms, indicating that most exchange events in *him-6* animals result from meiotic DSBs. Exchange chromosomes in *him-6* worms occured at a similar rate in both bivalents and univalents (22/46 and 4/14, respectively). Univalents are formed due to the failure of some inter-homolog crossover precursors to mature to chiasmata [40]. The pairwise comparisons between wildtype and *him-6* and between *him-6* and *him-6 spo-11* were significant (p<2.0001 and p<.00001, respectively; Pearson’s Chi-square test). N values indicate the number of chromatids counted. C. Mostly non-exchange univalents in a representative image of a *him-6 spo-11* diakinesis nucleus. Arrowhead indicates the single exchange chromosome. The two X chromosomes (asteriks) are unlabeled due to their late replication. Red, axis (anti-HTP-3 antibodies); green, EdU, Blue, DNA (DAPI). Scale bar = 1μm. D. Premeiotic and meiotic SCE formation in EdU labeled chromatids would result in indistinguishable diakinesis bivalents. Schematics of chromatids undergoing DNA replication with EdU, followed by pre-meiotic DNA replication without EdU, and undergoing SCEs (red asterisks) in meiosis (top) or during premeiotic DNA replication (bottom). Premeiotic SCEs could also occur at times other than premeiotic DNA replication (not shown).

### The Synaptonemal Complex promotes SCEs when it mislocalizes between sisters

Our observation that inter-sister crossovers are inhibited, directly or indirectly, by BLM^HIM-6^ raises the question of how inter-homolog crossovers are protected from BLM^HIM-6^ inhibition. Most inter-homolog crossovers are designated by an assemblage of pro-crossover factors – including COSA-1 and MutS◻ – that provide protection from disassembling activities [26,50]. The localization of these pro-crossover factors is directed by a conserved proteinaceous structure, the Synaptonemal Complex, that assembles exclusively at the interface between paired homologs [19,51–53].

In worms lacking the meiotic cohesin subunit REC-8, homologs mostly fail to pair and the Synaptonemal Complex is mislocalized to the inter-sister interface where it recruits procrossover factors [54,55]. These foci may represent inter-sister crossovers, suggesting that the Synaptonemal Complex can promote SCEs in *rec-8* animals in an analogous fashion to the promotion of inter-homolog crossovers in wildtype animals. EdU labeling in *rec-8* animals revealed elevated levels of SCEs (Fig. 4A and 4D), with about half of the chromosomes harboring an exchange. While this rate is significantly higher than in control animals, this analysis is confounded by chromosome fragmentation, by the poor univalent morphology in *rec-8* mutants, and the inability to distinguish SCEs based on axis localization in this background (Fig. 4A; [55]).

**Figure 4:**
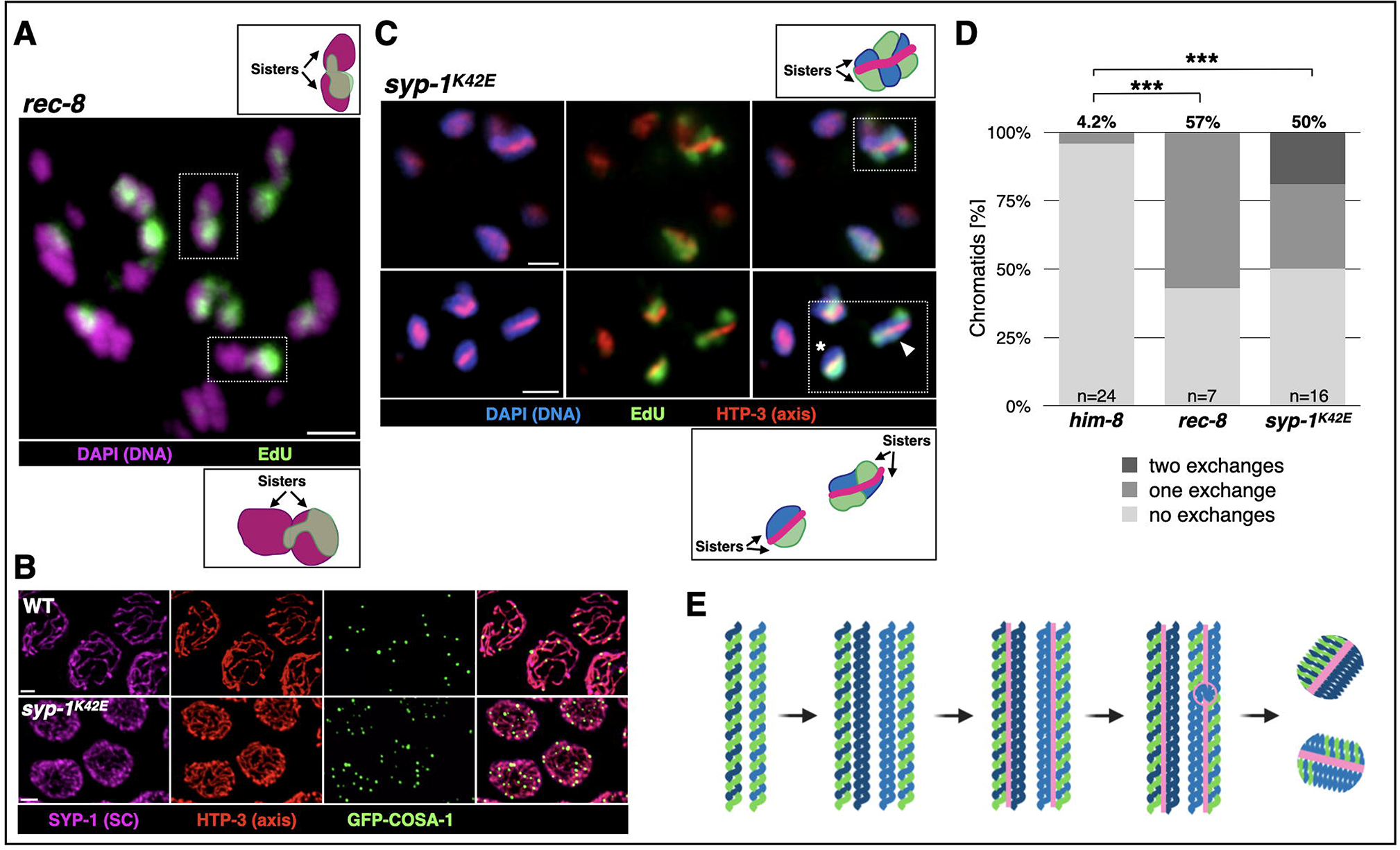
The Synaptonemal Complex promotes SCEs when it localizes between sister chromatids. A. Many exchange chromatids in an immunofluorescence image of a diakinesis nuclei from a *rec-8* animal. Purple, DNA (DAPI), green, EdU. Insets are explained in the interpretive diagrams above and below (boxed regions). Chromatids with crossovers are indicated by arrows. As observed previously [54,55,57], *rec-8* mutants affect chromosome morphology in diakinesis, and precluded the use of an axis marker to highlight SCEs. Scale bar = 1μm. B. The Synaptonemal Complex in *syp-1^K42E^* worms localizes to unpaired homologs but can recruit pro-crossover factors. Immunofluorescence image of pachytene nuclei from wildtype and *syp-1^K42E^* worms. The many more Synaptonemal Complex threads in *syp-1^K42E^* worms indicate localization to unpaired homologs. GFP-COSA-1 foci still localize to the Synaptonemal Complex. Purple, Synaptonemal Complex (SC; anti-SYP-1 antibodies); Red, axis (anti-HTP-3 antibodies); Green, GFP-COSA-1 (anti-GFP antibodies). Scale bars = 1μm. C. Univalent chromosomes undergoing multiple SCEs in immunofluorescence images of *syp-1^K42E^* worms grown at 25°C. Red, axis (anti-HTP-3 antibodies); green, EdU, Blue, chromatin (DAPI). Insets are explained in the interpretive diagrams above and below (boxed regions). Top, diakinesis nucleus containing a univalent with two SCEs. Bottom, a diakinesis nucleus highlighting a univalent with no exchanges (asterisk) and a univalent with a single SCE (arrowhead). Scale bar = 1μm. D. Elevated frequency of exchange univalents in *syp-1^K42E^* and *rec-8* worms. The unpaired X univalents in *him-8* animals (from Fig. 2A) are shown as controls. The percentages above the bars indicate the total number of exchange chromatids (p<2.01, for both pairwise comparisons to *him-8*, Pearson’s chi-square test; single and double SCEs were pooled together for statistical analysis). N values indicate the number of chromatids counted. E. A diagram illustrating Synaptonemal Complex localization between sisters in *rec-8* and *syp-1^K42E^* mutants, thereby promoting SCEs in an analogous way to the promotion of inter-homolog crossovers in wildtype animals.

We also observed elevated SCEs using an orthogonal perturbation: a mutation in the Synaptonemal Complex component *syp-1^K42E^* that mislocalizes the Synaptonemal Complex to the interface between sister chromatids, but does not impede the ability of the Synaptonemal Complex to recruit pro-crossover factors such as COSA-1 (Fig. 4B and SGG and OR, manuscript in preparation). We found elevated levels of SCEs in *syp-1^K42E^* animals (Fig. 4C-D), suggesting that pro-crossover factors now mark sites of future SCEs (Fig. 4E). Further supporting this idea, we observed both single and double SCEs in *syp-1^K42E^* animals, consistent with the presence of multiple GFP-COSA-1 foci per chromosome in this mutant (an average of 16.4 foci/nucleus; SGG and OR, manuscript in preparation).

Our analysis of *him-8* animals (Fig. 2) indicates that the Synaptonemal Complex does not prevent SCE formation, either directly or indirectly by bringing the homologs together. If that were the case, SCEs would have been elevated on the X chromosomes in *him-8* mutants, which are unpaired and not associated with the Synaptonemal Complex. We find, however, that the role of the Synaptonemal Complex in regulating crossovers extends beyond limiting the number of crossovers to exactly one per homolog pair [56]. It was previously shown that the Synaptonemal Complex promotes inter-homolog crossovers by recruitment of pro-crossover factors, thereby establishing a protective structure that prevents disassembly by helicases such as BLM^HIM-6^ [26,54,57]. Here we demonstrate that this idea also applies to inter-sister crossovers, which were likewise promoted by the Synaptonemal Complex when it is localized between the sisters (Fig. 4). Hence, by virtue of its localization exclusively between homologous chromosomes, the Synaptonemal Complex acts to limit crossover repair using the sister chromatid as a template (Fig. S3).

Our results also suggest that BLM^HIM-6^ disassembles repair intermediates that would otherwise mature to be SCEs, as was demonstrated for RecQ helicases in other systems ([44] and Fig. S3). Interestingly, removal of BLM^HIM-6^ does not increase the number of inter-homolog crossovers [40]. We hypothesize that underlying this seeming discrepancy is differential resolution: both inter-sister and inter-homolog intermediates form in *him-6* animals, but while inter-sister intermediates are resolved as both crossovers and noncrossovers, inter-homolog events preferentially yield noncrossovers. This idea is supported by the presence of transient inter-homolog associations in *him-6* mutants that lack bona fide crossovers [40]. It further suggests that a homolog-specific structure – potentially the Synaptonemal Complex or the axis – directs the resolution of repair intermediates.

## Conclusions

Using the cytological assay we developed, we show that sister-directed repair forms crossovers in <2% of cases. In contrast, the formation of the one inter-homolog crossover necessary to direct accurate chromosome segregation is promoted by a very robust assurance pathway [2,3]. Our analysis suggests that the low rate of SCEs is brought about by a dual-acting mechanism that designates and protects the single inter-homolog crossover, and that inhibits almost all other inter-homolog and inter-sister crossovers. While sister-directed repair is mostly error-free, DNA repair that involves repetitive elements can result in deletions, insertions and karyotype rearrangements. This risk is higher during crossover repair, since the integrity of both the broken chromosome and the repair template are compromised. We speculate that the meiotic program in worms goes to great lengths to limit crossover repair in order to mitigate the risk incurred by imperfect exchanges.

## STAR Methods

### Lead contact and materials availability

Requests for further information, reagents, or resources should be directed to the lead contact, Ofer Rog.

### Experimental model and subject details

Unless otherwise noted, *C. elegans* worms were cultured at 20°C under standard conditions [58]. *spo-11::aid(slc3)*, which degrades SPO-11 in the presence of auxin, was constructed using CRISPR/Cas9 RNP injection as described in [59], except without the 3×FLAG tag. Injection was performed into *ieSi38 [Psun-1::TIR1::mRuby::sun-1 3’UTR, cb-unc-119(+)] him-8 (tm611) IV* animals, resulting in linkage of all three elements. When grown on auxin, this strain is referred to in the text as *him-8 spo-11(−)* for simplicity. *syp-1^K42E^(slc11)* was obtained in a target mutagenesis screen, to be described elsewhere (SGG and OR, manuscript in preparation). *syp-1^K42E^* worms are temperature sensitive; they were maintained at 15°C, but were grown at 25°C prior to and during EdU labeling, causing the Synaptonemal Complex to localize to 12 unpaired homologs (Fig. 4B).

## Method details

### EdU incorporation

Worms were synchronized through egg washing as in [60]. Age matched worms were grown to L4 stage and then washed with 1ml of PBS with 0.1% Triton X (v/v) into a conical tube. Animals were washed one additional time to remove excess bacteria, and transferred to a new tube to which EdU (ThermoFisher A10044) dissolved in water as a 10mM stock was added to a final concentration of 4mM. Animals were then transferred to a nutator set to 80rpm and soaked in EdU for 40 minutes at room temperature. Finally, animals were washed twice with PBS/Triton X and plated onto fresh NGM plates. Once dry, the plates were then transferred to the 25°C incubator to begin an approximately 28 hour chase. Chase times were highly sensitive to genetic background and growth conditions, and were empirically determined to maximize single-labeled chromatids in diakinesis.

### Cytology

Cytology was performed as previously described in [61], with modifications to accommodate the click chemistry required for EdU visualization (ThermoFisher C10337). In brief, samples were fixed, dissected, and labeled with primary and secondary antibodies as in [61]. Slides were then washed for 30 minutes at room temperature in PBS with 1% Triton X (v/v) for further permeabilization to allow for better penetration of the ‘click’ chemistry reagents. Samples were then labeled with Alexa 488-azide according to the kit’s instructions for 30 minutes at room temperature. Samples were washed 3x 10 minutes and then DAPI stained for 20 minutes at room temperature. Slides were mounted using Prolong Glass (ThermoFisher P36980), cured overnight, and sealed with nail polish. In typical experiments, >95% of gonads exhibited bright EdU staining in meiotic prophase. The following antibodies were used: guinea-pig anti-HTP-3 (1:500, [62]), mouse anti-GFP (1:1000, Roche 11814460001), goat anti-SYP-1 (1:500, [63]), Cy3 AffiniPure Donkey Anti-Guinea pig (1:500, Jackson Immunoresearch), Cy5 AffiniPure Donkey anti-Goat (1:500, Jackson Immunoresearch), and 488 AffiniPure Donkey anti-Mouse (1:500, Jackson Immunoresearch).

### Depletion of proteins using auxin

NGM plates with a final concentration of 1μM auxin (indole-3-acetic acid, VWR AAA10556-36) were made according to [60]. P_0_ worms were grown until 24 hours post-L4 on standard NGM plates and then transferred to auxin plates to lay eggs. After 24 hours, the adult worms were removed and the F_1_ generation, whose full development occurred on auxin, were used as experimental animals.

### Irradiation

Worms were exposed to 5,000 rad (50 Gy) of X-ray irradiation using a W-(ISOTOPE) source, inducing an average of 7 DSBs per chromosome (14 per homolog pair). The number of DSBs generated per a given dose of radiation was empirically assessed by titrating low doses of radiation, as in [3], yielding a factor of 0.0028 DSBs per homolog pair per rad (SGG and OR, manuscript in preparation). Irradiation was performed either 1 hour or 6 hours after EdU labeling, for mitotic and meiotic induced breaks, respectively. Dissections were completed as described above, 28 hours after EdU labeling. For meiotic irradiation experiments (Fig. 2E), only nuclei with 7 DAPI staining bodies (5 bivalent autosomes and 2 X chromosome univalents) were considered for the analysis.

### Image acquisition and processing

Confocal Microscopy images were collected as z-stacks (at 0.2μm intervals) using a 63× 1.40 NA objective on a Zeiss LSM 880 microscope equipped with an AiryScan detector. Image processing and analysis was conducted using the ZEN software package (Blue 2.1). Partial maximum intensity projections are shown throughout.

### Quantification of exchanges

Exchanges between sisters were quantified at the diakinesis stage of meiotic prophase, where chromosomes are condensed and each sister chromatid is individualized. Depending on the presence or absence of a chiasma, the homologs condense in a bivalent cruciform structure or as two oval univalents, respectively (Figs. 1B and 2B). Several factors contributed to asynchrony in obtaining single-labeled sister chromatids in diakinesis: asynchronous mitotic cell-cycle, late replication of the X chromosome [64], and semi-synchronous entry into, and passage through, meiosis [2,11,12]. In addition, the poor resolution in the z-axis prohibited analysis of chromosomes that were not oriented horizontally relative to the imaging plane. Exchanges (or lack thereof) were examined only in chromosomes where one of the two sisters was labeled with EdU and the two sisters were clearly distinguishable in the xy-plane. Sister exchanges were scored when EdU labeled chromatin crossed the chromosome axis (decorated with anti-HTP-3 antibodies), which represents the inter-sister interface. For bivalents, each of the long arms was scored independently. In *rec-8* animals, the sisters were identified based on chromatin morphology.

KEY RESOURCES TABLE

KEY RESOURCES TABLE
| REAGENT or RESOURCE | SOURCE | IDENTIFIER |
| --- | --- | --- |
| **Antibodies** |  |  |
| Guinea pig anti-HTP-3 | Kim Lab | N/A |
| Mouse anti-GFP | Roche | Cat#11814460001 |
| Goat anti-SYP-1 | Dernburg Lab | N/A |
| Cy3 AffinPure Donkey anti-Guinea pig | Jackson ImmunoResearch | Cat#706-165-148 |
| Cy5 AffinPure Donkey anti-Goat | Jackson ImmunoResearch | Cat#705-175-147 |
| 488 AffiniPure Donkey anti-Mouse | Jackson ImmunoResearch | Cat#715-545-150 |
| **Chemicals, Peptides, and Recombinant Proteins** |  |  |
| Indole-3-acetic acid (auxin) | VWR | Cat#AAA10556-36 |
| Cas9 protein | IDT |  |
| EdU | ThermoFisher | Cat#A10044 |
| Prolong Glass antifade agent | ThermoFisher | Cat#P36980 |
| **Critical Commercial Assays** |  |  |
| Click-it EdU Cell Proliferation Kit | ThermoFisher | Cat#C10337 |
| **Experimental Models: Organisms/Strains** |  |  |
| *C. elegans*: N2 | CGC | N2 |
| *him-8(tm611)* IV | CGC | CA257 |
| *unc-119(ed3) III; ieSi38 [P<sub>sun-1</sub>::TIR1::mRuby::sun-1 3'UTR, cb-unc-119(+)]* IV | CGC | CA1199 |
| *lig-4(ok716) III* | CGC | RB873 |
| *him-6(ok412) IV* | CGC | VC193 |
| *rec-8(ok978) IV/nT1 [qls51] (IV;V)* | CGC | VC666 |
| *him-6(ok412) spo-11(me44) IV/nT1 (IV;V)* | CGC | TG2196 |
| *mels8 [pie-1p::GFP::cosa-1 + unc-119(+)] II* | CGC | AV620 |
| *unc-119(ed3) III; ieSi38 [P<sub>sun-1</sub>::TIR1::mRuby::sun-1 3'UTR, cb-unc-119(+)]* *him-8(tm611)* IV | This study | ROG125 |
| *unc-119(ed3) III; ieSi38 [P<sub>sun-1</sub>::TIR1::mRuby::sun-1 3'UTR, cb-unc-119(+)]* *him-8(tm611) spo-11::aid (slc3)* IV | This study | ROG130 |
| *syp-1<sup>K42E</sup>(slc11)* V | Described elsewhere (SGG and OR, manuscript in preparation) | ROG198 |
| *lig-4(ok717) III; him-8(tm611) IV* | This study | ROG189 |
| *mels8 [pie-1p::GFP::cosa-1 + unc-119(+)] II; syp-1<sup>K42E</sup>(slc11) V* | This study | ROG202 |
| **Software and Algorithms** |  |  |
| Zen 2.3 Blue Edition | Carl Zeiss Microscopy | N/A |
| Zen 2.3 Black Edition | Carl Zeiss Microscopy | N/A |
| RStudio Version 1.1.463 | RStudio, INC | N/A |
| **Other** |  |  |
| W-(ISOTOPE) X-ray source | Golic Lab | N/A |
| LSM880 confocal microscope with AiryScan | Carl Zeiss Microscopy | N/A |

## Quantification and statistical analysis

### Statistics

Chi-squared statistical analysis was conducted using the R software package. For pairwise comparisons of SCE rates (Figs. 2A, 3B and 4D), Pearson’s Chi Squared test was used. Our null hypothesis was that there is no change in SCE frequency, and a P value smaller than 0.05 was considered statistically significant and sufficient to reject the null hypothesis. In Figure S2A, an unpaired Student’s t-test was used. Our null hypothesis was that there was no difference between the two categories (with and without EdU), and a P value smaller than 0.05 was considered sufficient to reject the null hypothesis. Significance is marked throughout the figures using three asterisks above significant results. Confidence intervals shown in Fig. 2C-E represent 95% exact binomial confidence intervals.

## Acknowledgements

We would like to thank members of the Rog Lab for help with the cytology and statistical analysis and for critical reading of this manuscript; Tess Stapleton for her help with the R software package and with statistical analysis; Kent Golic for his feedback and for use of the X-ray source; Erik Toraason and Diana Libuda for sharing their manuscript and data prior to publication; Yuval Mazor, Yumi Kim, Keith Cheveralls and Sarit Smolikove for comments; Sara Nakielny for comments on the manuscript and editorial work; and Yumi Kim and Abby Dernburg for antibodies. Worm strains were provided by the CGC, which is funded by NIH Office of Research Infrastructure Programs (P40 OD010440). This work was supported by a Genetics Training Grant 5T32GM007464-42 to DEA, and by a Pilot Project Award from the American Cancer Society, 1R35GM128804 grant from NIGMS, and start-up funds from the University of Utah to OR.

## Authors contributions

DEA performed all the experimental work. SGG contributed reagents. DEA and OR conceptualized the project, analyzed the data, and wrote the manuscript.

---

**Supplementary Figure 1:**
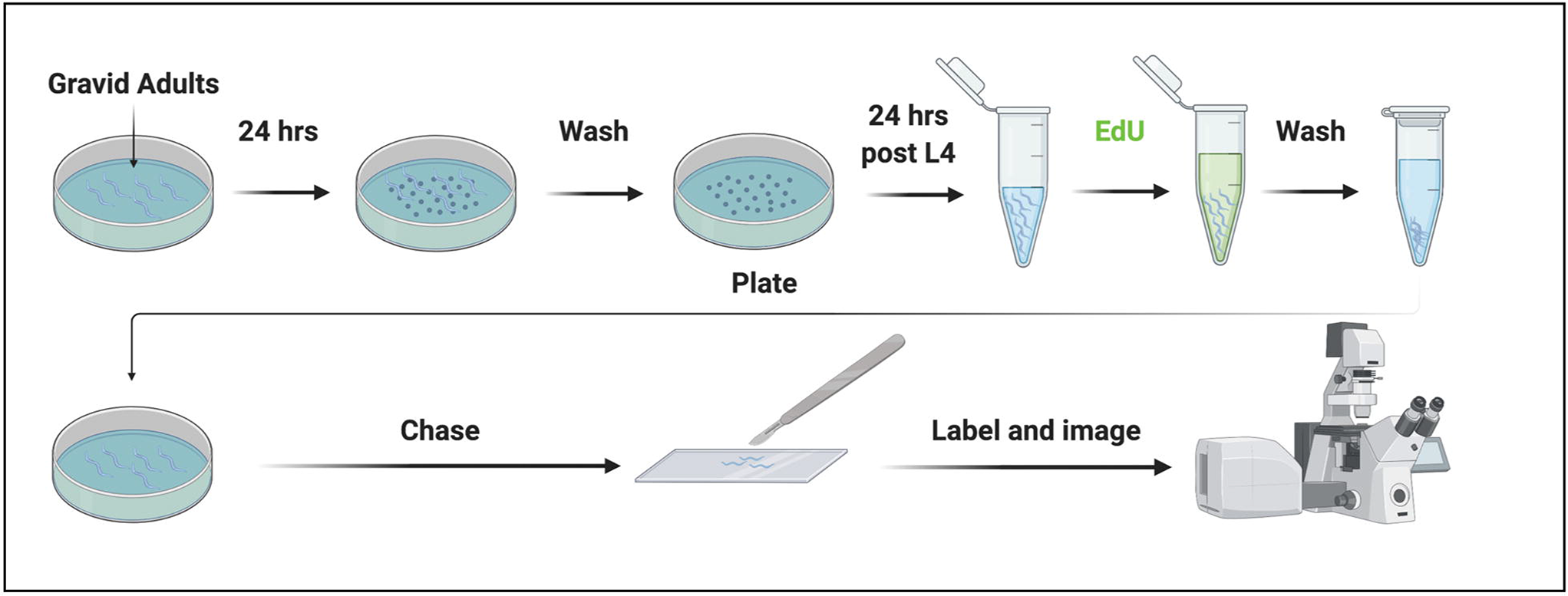
Graphical protocol for selective labeling with EdU. Experimental overview for EdU labeling. Gravid adult worms were transferred to a fresh plate to lay eggs. Adults were then removed by washing, and the eggs were grown to the pre-adult (L4) stage. Animals were then washed and incubated with EdU for 40 minutes. Following washes, worms were grown until single-sister labeled chromosomes reached diakinesis, when worms were dissected, fixed, stained with antibodies and with click chemistry to attach a fluorescent dye to EdU, and visualized by confocal microscopy.

**Supplementary Figure 2:**
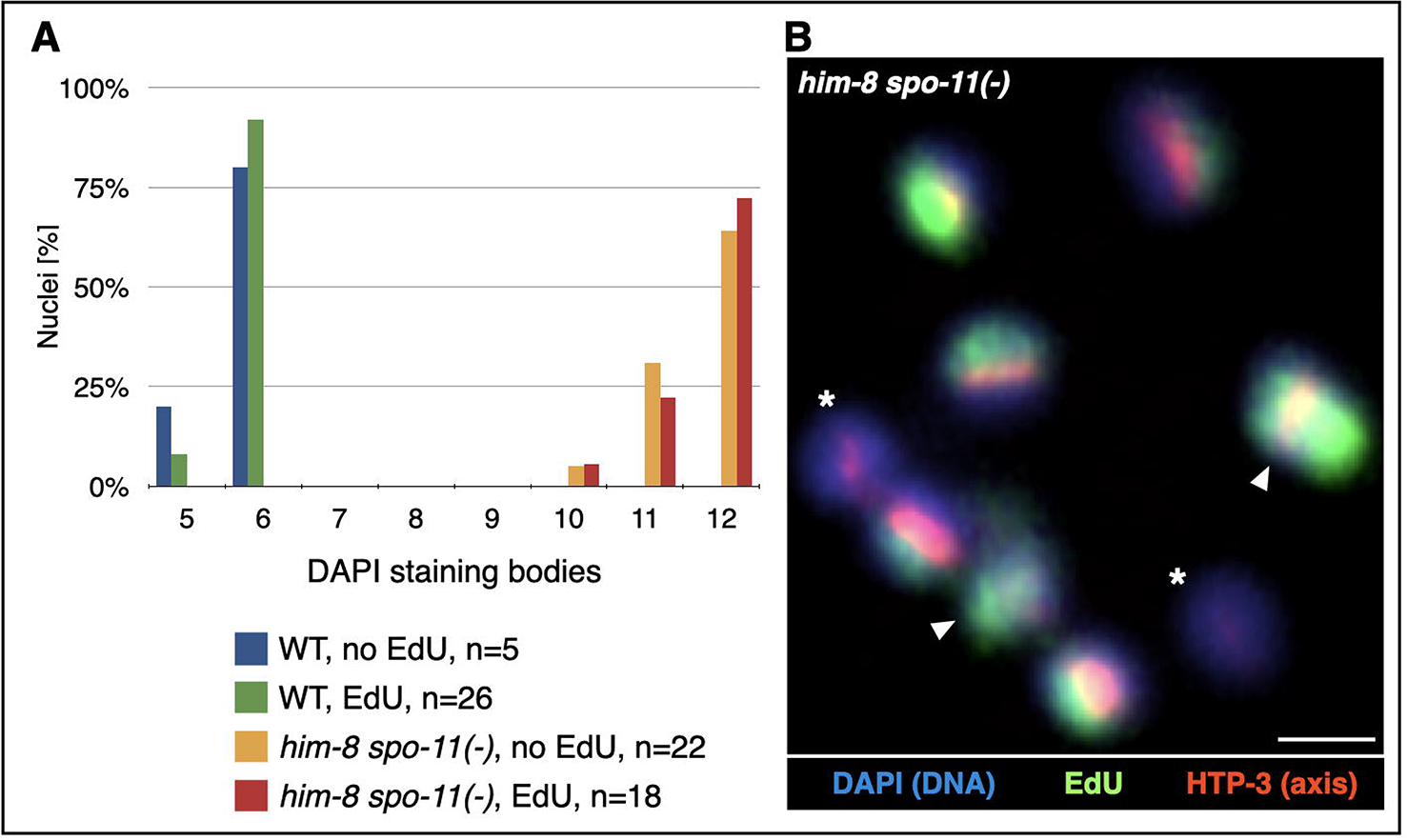
Further analysis of potential effects of EdU labeling. A. Analysis of chromosome masses in diakinesis (DAPI staining bodies) in wildtype and in *him-8 spo-11(−)* worms that lack meiotic DSBs. DAPI staining bodies are not significantly different between EdU-labeled and unlabeled chromosomes (unpaired Student’s t-test). The deviation from the expected 6 or 12 DAPI staining bodies (for wildtype or *him-8 spo-11(−)* animals, respectively) is explained by overlapping chromosomes. N values indicate the number of diakinesis nuclei counted. B. Lack of bivalents despite extensive EdU incorporation in a representative image of EdU labeled univalents in *him-8 spo-11(−)* animals. Arrowheads, overlapping chromosomes creating the appearance of two labeled sisters; asteriks, unlabeled X chromosomes as a result of differential DNA replication timing. Blue, DNA (DAPI); Green, EdU; Red, axis (anti-HTP-3 antibodies). Scale bar = 1μm.

**Supplementary Figure 3:**
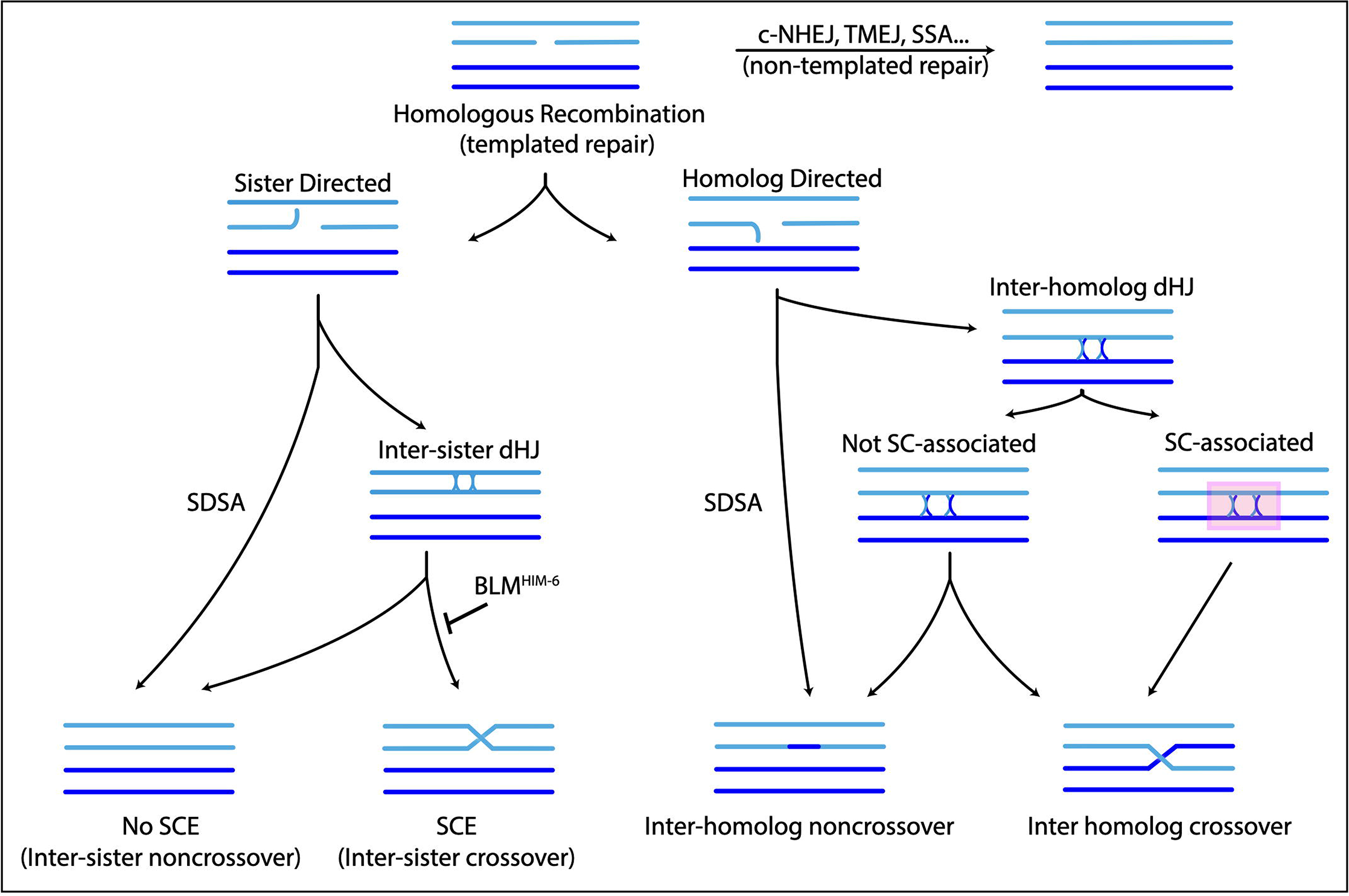
Model. Simplified model of meiotic break repair. DSBs are first segregated into templated, homologous recombination repair (bottom), and a disfavored non-templated repair (right) by mechanisms such as canonical non-homologous end-joining (c-NHEJ), theta-mediated end-joining (TMEJ), and single-strand annealing (SSA). Homologous recombination can use the sister chromatid as template (bottom left), or the homolog as a template (bottom middle). Double Holliday Junction (dHJ) intermediates in either case could be resolved as crossovers or noncrossovers. Noncrossovers could also form without dHJ intermediates, through synthesis-dependent strand annealing (SDSA). Homolog-directed crossover repair can occur in the context of the Synaptonemal Complex (SC) and of pro-crossover factors (bottom right), ensuring it will result in an inter-homolog crossover. Sister-directed repair rarely forms SCEs, partly due to inhibition by BLM^HIM-6^.

**Table S1:** Summary of all SCE counts.

| Genotype | Bivalents |  |  |  |  | Univalents |  |  |  |  |
| --- | --- | --- | --- | --- | --- | --- | --- | --- | --- | --- |
|  | Total | Exchanges | Non-Exchanges | % exchange chromatids (±95% CI) | % SCEs/DSB (±95% CI)<sup>a</sup> | Total | Exchanges | Non-exchanges | % exchange chromatids (±95% CI) | % SCEs/DSB (±95% CI)<sup>a</sup> |
| WT | 49 | 2 | 47 | 4.08%<br>(0.5%-13.98%) | 1.63%<br>(0.20%-5.59%) | - | - | - | - | - |
| *him-8* | 35 | 0 | 35 | 0%<br>(0%-10%) | 0%<br>(0%-4.00%) | 24 | 1 | 23 | 4.17%<br>(0.11%-21.12%) | 1.67%<br>(0.04%-8.45%) |
| *lig-4; him-8* | 55 | 1 | 54 | 1.82%<br>(0.05%-9.72%) | 0.73%<br>(0.02%-3.89%) | 28 | 0 | 28 | 0%<br>(0%-12.34%) | 0% (0%-4.94%) |
| *him-8 spo-11(-)*<sup>b</sup> | - | - | - | - | - | 30 | 1 | 29 | 3.33%<br>(0.08%-17.22%) | NA |
| *him-8 spo-11(-) + mitotic IR* | - | - | - | - | - | 15 | 2 | 13 | 13.33%<br>(1.66%-40.46%) | 1.90%<br>(0.22%-5.39%) |
| *him-8 spo-11(-) + meiotic IR* | 20 | 2 | 18 | 10%<br>(1.24%-31.70%) | 1.43%<br>(0.17%-4.23%) | 4<sup>c</sup> | 2 | 2 | NA | NA |
| *rec-8* | - | - | - | - | - | 7 | 4 | 3 | 57.14%<br>(18.41%-90.10%) | 22.86%<br>(7.36%-36.04%) |
| *syp-1<sup>K42E</sup>* | - | - | - | - | - | 16 | 8 | 8 | 50%<br>(24.65%-75.35%) | 20%<br>(9.86%-30.14%) |
| *him-6<sup>d</sup>* | 46 | 22 | 24 | 47.83%<br>(32.89%-63.05%) | 19.13%<br>(13.16%-25.22%) | 14 | 4 | 10 | 28.57%<br>(8.39%-58.10%) | 11.43%<br>(3.36%-23.24%) |
| *him-6 spo-11* | - | - | - | - | - | 49 | 6 | 43 | 12.24%<br>(4.63%-24.77%) | 4.90%<br>(1.85%-9.91%) |

<sup>a</sup> The ratio was calculated based on the assumption of 2.5 DSB per chromosome, except in the case of irradiated animals, where the measured average of 7 DSB per chromosome was used.
<sup>b</sup> There was no statistically significant difference between *him-8* and *him-8 spo-11(-)* animals that lack meiotic DSBs. Therefore, while the SCEs we observed in WT, *him-8* and *lig-4; him-8* animals are likely an outcome of repairing meiotically induced DSBs, their low frequency prevented us differentiating them from events occurring during DNA replication.
<sup>c</sup> The univalent chromosome were the unpaired X chromosomes. Their small number precluded statistical analysis.
<sup>d</sup> Bivalents and univalents were pooled together in Fig. 3B. All the chromosomes are paired and synapsed in *him-6* mutants, and harbor one GFP-COSA-1 focus. The lack of chiasmata on some chromosomes reflects a role for BLM<sup>HIM-6</sup> in ensuring the maturation of these precursors to chiasmata.

